# An interpretable molecular framework for predicting cancer driver missense mutations

**DOI:** 10.1101/2025.09.03.674130

**Authors:** Yan Yang, Weikang Sun, Jian Zhang, Yang Liu, Minghui Li

## Abstract

Missense mutations play a critical role in human disease, contributing to both inherited disorders and cancer. However, accurately predicting their functional impact—particularly for cancer driver mutations—remains a major challenge due to limited validated labels and the complex molecular basis of oncogenesis. Here, we systematically characterized over 120,000 missense variants across pathogenic, benign, driver, passenger, recurrent somatic, and common population classes, using a comprehensive set of mechanistically grounded molecular features. By assessing the statistical burden of variations, we demonstrated that these features effectively discriminate among diverse variant classes and reveal a consistent enrichment of functional sites, structural integrity, and biophysical changes in pathogenic and driver mutations. Building on these insights, we developed MutaPheno, an interpretable framework for predicting the functional consequences of missense mutations. The model integrates 34 molecular-level features, encompassing structural, functional, physicochemical, and contextual descriptors, using a random forest algorithm. Trained exclusively on pathogenic and benign variants, MutaPheno achieved strong accuracy in predicting cancer driver mutations, outperforming both cancer-specific and general pathogenicity tools, while also demonstrating superior robustness when tested on unseen proteins. Our findings highlight the shared mechanisms between pathogenic and driver mutations and emphasize the role of molecular features in improving variant interpretation. MutaPheno provides a transparent and generalizable tool that can facilitate driver discovery and the development of targeted therapies.

## Introduction

Missense mutations—single-nucleotide substitutions that result in amino acid changes—are among the most prevalent and functionally significant genetic variants. By altering protein structure, stability, or interaction networks, these mutations can disrupt essential biological processes and contribute to disease^1, 2^. When such mutations impair protein function and lead to disease, they are referred to as pathogenic missense mutations. Classic examples include the E6V substitution in the β-globin chain causing sickle cell disease^3^ and missense mutations in APP, PSEN1, and PSEN2 have been shown to be causative of early-onset familial Alzheimer’s disease^4^. Whereas hereditary disorders often result from germline variants, cancer is primarily driven by the accumulation of somatic mutations. Within the complex mutational landscape of tumors, only a small fraction—known as driver mutations—confer a selective growth advantage and actively promote tumorigenesis^5^ while the majority are passenger mutations with no functional consequence^6^. Missense mutations that promote oncogenesis are classified as cancer driver missense mutations. Representative examples of well-characterized driver mutations include the KRAS G12D substitution frequently observed in pancreatic ductal adenocarcinoma^7^, the TP53 R175H hotspot mutation identified across a broad spectrum of carcinomas^8^, and the BRAF V600E mutation, a key oncogenic event in melanoma^9^.

Understanding and accurately characterizing the functional impact of missense variants is essential for disease diagnosis and the development of targeted therapies. While experimental assays remain the gold standard for functional validation, their high cost and low throughput limit their scalability. Consequently, over 40 computational tools have been developed to predict the pathogenicity of missense mutations, drawing on features such as evolutionary conservation, protein structural properties, and machine learning algorithms^10, 11^. However, the vast majority of these tools were originally designed for inherited diseases. Representative examples—including SIFT^12^, PolyPhen-2^13^, REVEL^14^, VEST4^15^, and AlphaMissense^16^—have shown strong performance in identifying pathogenic variants. In contrast, only a limited number of tools have been specifically tailored for predicting cancer driver missense mutations, with notable examples such as CHASM^17^, ParsSNP^18^, CHASMplus^19^, and AI-Driver^20^ addressing cancer-specific challenges like somatic mutation patterns and tumor heterogeneity. Despite these advances, existing computational approaches still suffer from two major limitations: they often operate as “black boxes” with limited mechanistic interpretability, which hinders the development of targeted therapies; and they exhibit limited predictive accuracy and poor generalizability to cancer contexts—challenges that are further exacerbated by the scarcity of experimentally validated driver mutations and the high genetic heterogeneity of tumors^21-23^.

A promising strategy to overcome the limitations of existing variant effect predictors is to focus on molecular phenotypes—quantifiable alterations at the molecular level caused by missense mutations, such as changes in structural stability, disruption of interaction interfaces, or modification of post-translational modification sites^24^. While not all molecular phenotypes manifest as observable organismal traits, most phenotypic and fitness-related consequences ultimately stem from such molecular disruptions. Leveraging these mechanistic insights allows predictive models to achieve greater interpretability by incorporating molecular features. Over the past decade, a series of studies have explored how molecular features reveal the functional impact of missense mutations and help distinguish pathogenic, driver, and benign variants. In 2016, we showed that integrating multiple protein conformations with stability and binding affinity assessments effectively predicts the functional impact of cancer mutations in CBL^25^. Iqbal et al. identified physicochemical, structural, and functional differences between pathogenic and population variants, uncovering both shared and protein class–specific signatures^26^. Cheng et al. found that germline pathogenic variants are significantly enriched at protein–protein interaction interfaces^27^. Laddach et al. extended this line of work by analyzing variant enrichment across structural regions and functional domains, integrating multi-omics data to distinguish pathogenic, rare, and common variants^28^. Most recently, Gerasimavicius et al. showed that different mutation types—loss-of-function, gain-of-function, and dominant-negative—exhibit distinct structural patterns^29^.

Despite recent progress, key limitations remain. First, most studies focus on a narrow set of molecular-level features—such as interaction interfaces, post-translational modification sites, or protein stability—to identify statistical differences across mutation classes^25-30^. This limits their ability to comprehensively describe mutation effects across biological contexts and hinders a full mechanistic understanding. Second, many efforts remain descriptive and fail to translate molecular features into interpretable predictive models. Although some machine learning approaches incorporate molecular features, they often depend on external scores—such as pathogenicity annotations or conservation metrics—rather than building models from intrinsic, mechanistically meaningful descriptors^26^. This reliance ultimately limits both interpretability and generalizability.

To address these limitations, we developed MutaPheno, a unified and interpretable classifier that predicts the functional impact of missense mutations based on curated and computed molecular-level features. MutaPheno integrates 34 structural, functional, physicochemical, and contextual descriptors that systematically characterize pathogenic, benign, driver, passenger, recurrent somatic, and common population variants. These features capture shared and context-specific molecular signatures across variant classes. By training a random forest model exclusively on pathogenic and benign variants, MutaPheno achieves high accuracy and generalizability across both germline and somatic contexts. Notably, despite lacking driver annotations during training, it performs competitively in driver mutation prediction, underscoring the shared molecular basis of functional mutations across diseases. Our work highlights the value of biologically grounded, phenotype-driven models for robust, interpretable, and generalizable prediction of missense mutation effects, and provides a framework for future studies linking molecular mechanisms to disease phenotypes.

## Methods

### Data collection and processing for missense mutations

#### Pathogenic and benign mutations

We curated missense mutations from three sources: ClinVar^31^, UniProt Humsavar^32^, and the dataset compiled by Sevim Bayrak et al.^33^. Fig. 1a provides an overview of the data sources and mutation categories analyzed in this study, and detailed version information, release dates, and access links are listed in Table S1. Mutations from ClinVar and Humsavar were retained only if annotated as “Pathogenic” or “Likely pathogenic” and “Benign” or “Likely benign”, respectively; entries with conflicting interpretations were excluded. From the Sevim Bayrak dataset, we extracted gain-of-function (GOF) and loss-of-function (LOF) missense mutations, originally curated from the 2019 release of HGMD^34^. All variants labeled as “Pathogenic” or “Likely pathogenic,” together with the GOF and LOF variants, were categorized as pathogenic, while those labeled as “Benign” or “Likely benign” were classified as benign. The counts of processed variants are summarized in Table S2.

**Figure 1.**
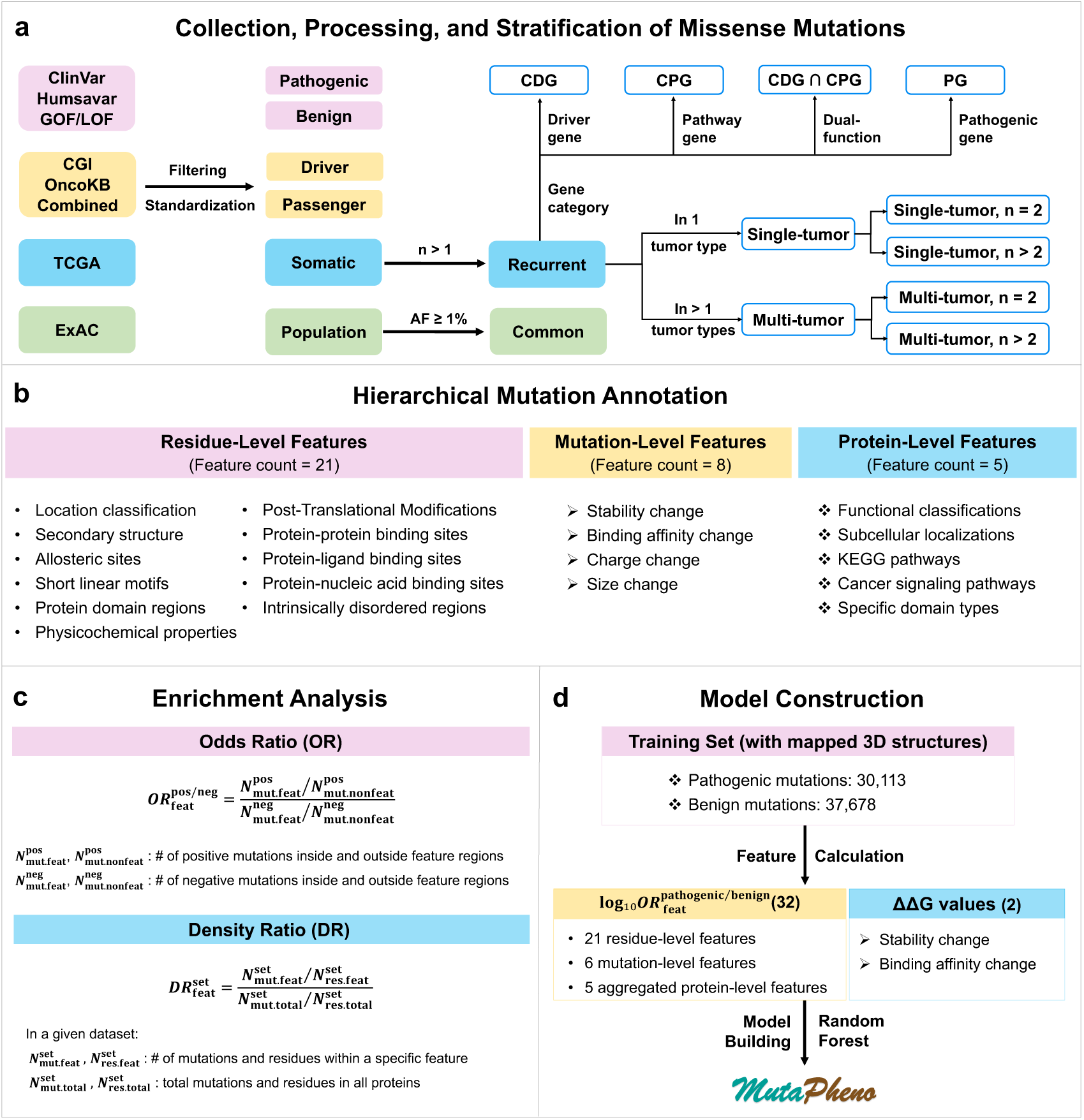
Overview of the study workflow. (a) Data collection, processing, and stratification of missense variants. Variants were categorized into six functional classes: pathogenic, benign, driver, passenger, recurrent somatic, and common population variants. Recurrent mutations were further stratified by tumor-type specificity and gene-level annotation. (b) Feature annotation. Each variant was annotated with 34 molecular features, including 21 residue-level, 8 mutation-level, and 5 protein-level features. (c) Statistical analysis. Feature-level associations were assessed using odds ratio (OR) for relative enrichment and density ratio (DR) for absolute enrichment. (d) Model construction. The MutaPheno random forest classifier was trained using a curated set of pathogenic and benign variants mapped to AlphaFold2 structures, incorporating 32 OR-weighted features and two ΔΔG-based structural perturbation features.

#### Driver and passenger mutations

Driver and passenger mutations were collected from OncoKB^35^, the Cancer Genome Interpreter (CGI)^36^, and our previously published dataset, referred to as the Combined Set^37^. From OncoKB, we included variants annotated as “Oncogenic,” “Likely Oncogenic,” or “Likely Neutral.” From CGI, we retained all validated oncogenic mutations. In our prior study^37^, we compiled experimentally characterized missense mutations across 58 cancer genes and classified them as “non-neutral” or “neutral” based on functional impact. For the current study, mutations labeled as “Oncogenic” or “Likely Oncogenic” in OncoKB, oncogenic variants from CGI, and “non-neutral” variants from the Combined Set were categorized as drivers. In contrast, mutations labeled as “Likely Neutral” in OncoKB and “neutral” in the Combined Set were categorized as passengers. Variant counts are listed in Table S2.

#### Somatic and population mutations

We collected somatic mutations from TCGA, as curated by Ellrott et al.^38^, and applied the filtering criteria of Bailey et al.^39^, yielding 746,392 missense mutations across 33 cancer types (Tables S2). Population variants were obtained from the non-TCGA subset of ExAC, as compiled in gnomAD database^40^, excluding mutations without a “PASS” filter. For comparative analyses, TCGA missense mutations were stratified by recurrence into Single (observed in one tumor sample, *n* = 1) and Recurrent (observed in multiple samples, *n* > 1), while ExAC variants were grouped by allele frequency (AF) into Rare (AF < 1%) and Common (AF ≥ 1%). Prior studies have shown that recurrent somatic mutations are more likely cancer drivers^41^, whereas common population variants are typically benign^40^. Based on this, we focused on comparing recurrent somatic mutations and common population variants to identify molecular features distinguishing likely drivers from frequently observed benign mutations.

### Standardization of mutations via identifier mapping and sequence alignment

To facilitate downstream analysis, all proteins and their associated mutations—originally annotated using transcript-, protein-, or gene-level identifiers from various data sources (Table S2)—were standardized to UniProt canonical accessions with matched amino acid substitutions. This step was essential, as subsequent annotations were performed at the protein sequence and structure level. RefSeq transcript IDs, and Ensembl transcript/protein IDs were mapped to UniProt isoform accessions via the EBI Proteins API^42^, while gene symbols were converted using the UniProt ID mapping tool^43^. Isoform-specific mutations were aligned to canonical sequences using Clustal Omega (v1.2.4)^44^ to determine accurate residue positions. To reduce potential bias in downstream statistical analyses, proteins longer than 3,000 amino acids (approximately 0.82% of the dataset) were excluded. The final counts of proteins and mutations retained for analysis are presented in Table 1, while the corresponding counts mapped to AlphaFold2 structures are shown in Table S3.

**Table 1.** Overview of mutation datasets analyzed in this study. This table presents the number of mutations, unique mutated sites, and corresponding proteins across diverse variant categories.

| **Dataset** | **# of mutations** | **# of sites** | **# of proteins** |
| --- | --- | --- | --- |
| Pathogenic | 36,613 | 31,044 | 3,206 |
| Benign | 51,844 | 50,942 | 12,744 |
| Driver | 3,720 | 2,497 | 262 |
| Passenger | 4,556 | 3,171 | 108 |
| Recurrent | 24,080 | 23,534 | 8,925 |
| Common | 24,621 | 24,406 | 9,713 |
| *Recurrent variants were classified based on tumor-type specificity* |  |  |  |
| Single-tumor | 6,624 | 6,591 | 3,839 |
| Multi-tumor | 17,456 | 17,025 | 7,775 |
| Single-tumor, n=2 | 5,844 | 5,828 | 3,647 |
| Multi-tumor, n=2 | 15,072 | 14,971 | 7,417 |
| Single-tumor, n>2 | 780 | 775 | 601 |
| Multi-tumor, n>2 | 2,384 | 2,177 | 1,515 |
| *Recurrent variants were classified based on gene-level annotation* |  |  |  |
| CDG | 1,905 | 1,829 | 475 |
| CPG | 953 | 948 | 387 |
| CDG∩CPG | 1,195 | 903 | 184 |
| PG | 1,360 | 1,350 | 451 |
| Total | 127,176 | 117,042 | 15,758 |

### Stratification of recurrent somatic mutations by tumor type and gene category

To assess differences in oncogenic potential, recurrent variants were first classified as single-tumor (recurrent in one cancer type) or multi-tumor (shared across multiple cancer types), and further stratified by recurrence frequency into four subgroups: single-tumor (*n* = 2), single-tumor (*n* > 2), multi-tumor (*n* = 2), and multi-tumor (*n* > 2) (Fig. 1a). This stratification enabled controlled comparisons of mutation enrichment patterns by recurrence level and tumor-type specificity.

To explore differences across gene categories, we curated high-confidence cancer driver genes (CDG), cancer pathway genes (CPG), and non-cancer pathogenic genes (PG). The CDG set was integrated from IntOGen^45^, OncoVar^46^, NCG^47^, Cancer Gene Census^48^, and ClinGen^49^, yielding 1,050 unique genes. CPG were compiled from 15 cancer-related KEGG pathways in MSigDB^50^ and 32 curated signaling pathways in NetSlim^51^, comprising 1,042 genes. PG were obtained from ClinGen^49^ by selecting genes with “Definitive” clinical validity and no cancer-related phenotype, resulting in 1,117 non-cancer disease genes.

To enable mutually exclusive comparisons, 272 proteins overlapping between CDG and CPG were defined as dual-function genes (CDG∩CPG). After removing these, 786 CDG-specific and 782 CPG-specific proteins remained. From the PG set, genes overlapping with cancer-related categories were excluded, yielding 922 proteins exclusively associated with non-cancer diseases. Recurrent mutations were mapped to one of the four gene classes—CDG, CPG, CDG∩CPG, or PG—based on their protein annotations (Fig. 1a).

### Hierarchical annotation of missense mutations at the residue and mutation levels

We annotated missense mutations using a hierarchical framework that integrates residue-level, mutation-level, and protein-level features (Fig. 1b). Residue-level features describe the structural and functional environment of the mutated site, including local geometry, binding interfaces, etc. Mutation-level features characterize the specific physicochemical changes introduced by amino acid substitutions, such as alterations in stability and binding affinity. Protein-level features reflect broader biological properties of the affected protein and provide context for interpreting the mutation’s potential impact. In this section, we provide a brief overview of the first two feature types; detailed definitions, methods, and statistics are provided in the Supplementary Information.

Residue-level features:

- Location classification: Residues were classified as buried (Core; rSASA < 0.2) or surface-exposed (Surface) based on relative solvent-accessible surface area (rSASA), calculated using DSSP^52^.
- Secondary structure: Residues were assigned to Helix, Sheet, or Loop categories based on DSSP annotations^52^.
- Allosteric sites (AlloSite): Combined experimentally validated sites from the Allosteric Database (ASD)^53^ and predicted sites from AlloSitePro^54^.
- Protein–protein binding sites (PPI-BS): Integrated experimentally determined interfaces from PDB crystal structures and predicted binding sites from Interactome INSIDER^55^.
- Protein–ligand binding sites (PLI-BS): Derived from ligand-bound PDB structures and predicted using P2Rank^56^.
- Protein–nucleic acid binding sites (PNI-BS): Extracted from PDB structures and expanded using GraphBind^57^ predictions for 2,047 nucleic acid-binding proteins identified from ENPD^58^ and EuRBPDB^59^.
- Post-translational modifications (PTM): Collected from PhosphoSitePlus^60^ and dbPTM^61^; residues located within 4 Å and 8 Å of PTM sites, including the modified residues themselves, were labeled PTM-4Å and PTM-8Å, respectively.
- Intrinsically disordered regions (IDReg): Verified disordered segments obtained from MobiDB^62^ and DisProt^63^.
- Short linear motifs (SLiM): Sourced from the ELM database^64^.
- Protein domain regions (DomReg): Domain annotations retrieved from InterPro^65^; residues within annotated domains were labeled DomReg.
- Physicochemical properties: The 20 standard amino acids were grouped into six categories: aliphatic (Ala, Ile, Leu, Met, Val), aromatic (Phe, Trp, Tyr), positively charged (His, Lys, Arg), negatively charged (Asp, Glu), neutral (Asn, Gln, Ser, Thr), and special (Cys, Pro, Gly).

Mutation-level features:

- Stability change (ΔΔ*G*_*fold*_): Predicted using PremPS^66^, developed by our group, based on AlphaFold2 structures; positive values indicate destabilization, negative values stabilization.
- Binding affinity change (ΔΔ*G*_*bind*_): Predicted using MutaBind2^67^, developed by our group, based on experimentally resolved heterodimer structures; positive values indicate weakened binding, negative values strengthened binding.
- Charge change: Amino acids assigned integer charges (+1: Lys, Arg; –1: Asp, Glu; 0: others); mutations categorized as Charge-0 (no change), Charge-1 (moderate change), or Charge-2 (strong change) based on absolute difference.
- Size change: Amino acids grouped as small (0: Ala, Gly, Ser, Cys, Pro, Thr, Asp, Asn), medium (1: Val, His, Glu, Gln), or large (2: Ile, Leu, Met, Lys, Arg, Phe, Trp, Tyr); mutations classified as Size-0, Size-1, or Size-2 by size difference.

In total, we compiled and computed 21 residue-level features: 11 structure-based (Core, Surface, Helix, Sheet, Loop, AlloSite, PPI-BS, PLI-BS, PNI-BS, PTM-4Å, PTM-8Å) and 10 sequence-based (PTM, IDReg, SLiM, DomReg, Aliphatic, Aromatic, PosCharge, NegCharge, Neutral, Special). Among the structure-based features, annotations for allosteric sites and predicted PPI-BS were directly obtained from external databases, while the others were derived through in-house computations using experimentally resolved structures from the Protein Data Bank (PDB)^68^ and/or AlphaFold2-predicted monomeric models^69, 70^. Models that were truncated or had a median pLDDT score below 70 were excluded. Residue-level features were assigned to each mutation based on the properties of the mutated site. In addition, we defined eight mutation-level features: two structure-based descriptors (ΔΔ*G*_*fold*_ and ΔΔ*G*_*bind*_) and six sequence-based indicators reflecting changes in charge and size (Charge-0/1/2 and Size-0/1/2).

### Odds ratio analysis of feature enrichment in positive versus negative mutations

The odds ratio (OR) is a statistical metric used to quantify the relative likelihood of an event occurring in one group compared to another^26, 29^. We calculated ORs to evaluate whether positive mutations (e.g., pathogenic, driver) are more likely than negative mutations (e.g., benign, passenger) to be enriched in specific categories of 21 residue-level and six mutation-level features (including charge and size changes) (Fig. 1c). ORs were calculated as:

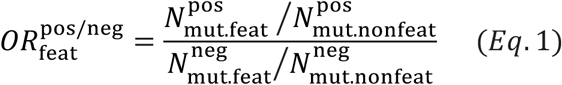

Where 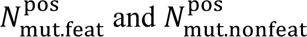 are the counts of positive mutations within or outside the feature region, respectively, and 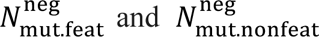 are the corresponding counts for negative mutations. For example, 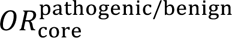 quantifies whether pathogenic mutations are more likely than benign mutations to occur in the core region. Details for this example are provided in the Supplementary Information. For sequence-based features, the entire protein sequence was considered; for structure-based features, only structurally resolved regions were included. Proteins lacking relevant feature coverage were excluded.

ORs were computed using two-sided Fisher’s exact tests implemented in the SciPy (Python)^71^. P-values were adjusted for multiple testing using the Benjamini–Hochberg (BH) procedure^72^ to obtain q-values. The 95% confidence intervals (CIs) were estimated using the Wald method^73^. A feature was considered significantly associated with mutation status if OR ≠ 1, the 95% CI did not include 1, and q < 0.05. For visualization, log₁₀OR values were used: log₁₀OR > 0 indicates enrichment in positive mutations, while log₁₀OR < 0 indicates enrichment in negative mutations.

### Variant set-specific feature enrichment analysis

Although odds ratios (ORs) effectively compare the relative prevalence of positive versus negative mutations within a given feature, they do not measure the absolute enrichment of mutations in that feature relative to overall dataset mutation density. To address this limitation, we employed the Density Ratio (DR) metric^28, 74^, defined in Equation 2:

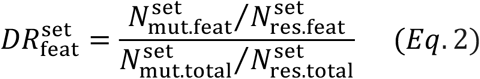

Here, 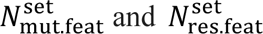 represent the number of mutations and residues within a given feature, respectively, for a specific dataset. 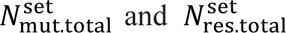 are the total number of mutations and residues across all analyzed proteins. For example, 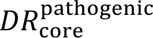 quantifies the density of pathogenic mutations in core regions relative to their overall density in the pathogenic dataset. Details for this example are provided in the Supplementary Information.

To assess statistical significance, we generated a null distribution of DR values from 2,000 random simulations per dataset, randomly reassigning mutations to positions within the same protein while preserving mutation counts per protein and allowing multiple mutations at the same residue. Two-tailed p-values were obtained by comparing observed DR values to this null distribution. P-values were adjusted for multiple testing using the BH method to obtain q-values. DR values were transformed to log₁₀DR for visualization; log₁₀DR > 0 indicates feature enrichment. To estimate 95% confidence intervals (CIs), we performed 2,000 bootstrap resamplings with replacement at the mutation level, deriving CIs from the 2.5th and 97.5th percentiles of the bootstrap distribution.

### Integrative protein-level features for mutation impact modeling

We annotated proteins using five protein-level features: functional classes, subcellular localizations, KEGG pathways, cancer signaling pathways, and protein domain types. Unlike residue- and mutation-level features, protein-level annotations were not analyzed individually for enrichment; they were used solely for model construction in MutaPheno. Because each protein-level annotation category contains many subclasses, analyzing them individually would result in very limited mutation counts per group. A brief overview is provided below, and detailed definitions and components of the five protein-level features are available in the Supplementary Information and Data S1.

- Functional classes were retrieved from the PANTHER classification system^75^, which assigns proteins to 24 groups based on conserved functions and evolutionary relationships.
- Subcellular localization data were obtained from the Human Protein Atlas (HPA)^76^, including only high-confidence annotations (Enhanced, Supported, or Approved) covering 36 compartments.
- Pathway annotations were sourced from 186 KEGG pathways in MSigDB^50^ and 32 curated cancer signaling pathways in NetSlim^51^.
- Protein domain types were annotated using InterPro^65^; subdomains were consolidated into parent categories, yielding 4,608 unique domain types.

To incorporate these annotations into modeling, we computed five aggregated protein-level features—one per annotation category. For each mutation, we identified all annotation terms within a category associated with its protein and summed the log-transformed odds ratios comparing pathogenic and benign mutations to obtain the aggregated score:

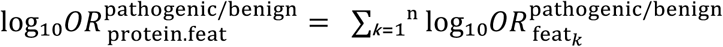

where *n* is the number of annotation terms in the category, and 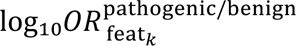 is the log-transformed odds ratio for the *k*-th term. The aggregated score reflects the cumulative pathogenic enrichment of a protein in each category. Additional details and illustrative examples are provided in the Supplementary Information.

### Construction of the MutaPheno model

#### Training set

We curated a high-quality dataset of pathogenic and benign missense mutations as described above (Fig. 1a and Table 1). To ensure the completeness and reliability of structure-based features, only variants mapped to AlphaFold2-predicted 3D structures with no truncation and a median pLDDT ≥ 70 were retained (Table S3). The final training set comprised 67,791 missense mutations from 9,928 proteins (30,113 pathogenic and 37,678 benign variants).

#### Independent test sets

To evaluate generalizability, we prepared two independent test sets:

- *Pathogenic mutation test set.* Derived from AlphaMissense^16^ and CPT-1^77^ test sets (ClinVar sourced), harmonized and filtered for canonical alignment and non-overlap with our training data. After mapping to AlphaFold2 structures, this set included 28,717 mutations (12,539 pathogenic, 16,178 benign).
- *Driver mutation test set.* Curated from the driver/passenger dataset described previously (Fig. 1a, Table 3). Variants overlapping with training data were excluded, yielding 6,322 mutations (2,317 drivers, 4,005 passengers).

#### Feature engineering

We incorporated 34 features: 21 residue-level, 8 mutation-level, and 5 aggregated protein-level annotations. For the residue-level features and six mutation-level features (charge and size changes), we calculated 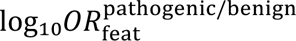 and encoded features as binary indicators multiplied by their log₁₀OR values, integrating feature presence and pathogenic enrichment. The two mutation-level biophysical features (folding stability and binding affinity changes) were represented by ΔΔG values from PremPS and MutaBind2; unavailable predictions were assigned a value of 0. For protein-level features, we used aggregated 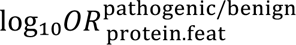 scores as described earlier.

#### Model construction and hyperparameter tuning

We constructed MutaPheno using a random forest (RF) classifier, optimizing two key hyperparameters: the number of trees (*n_estimators*, 200 to 2500 in steps of 50) and the maximum number of features considered at each split (*max_features*, set to 1 to 34). In our previous studies^66, 67, 78-80^, the RF algorithm was rigorously validated and consistently outperformed other conventional machine learning methods. Its advantages include high computational efficiency, robustness to overfitting, and the ability to generate interpretable feature importance scores.

To reduce sequence redundancy and improve model generalization, we partitioned the data based on protein sequence similarity. Pairwise sequence alignment was performed using MMseqs2^81^, with minimum thresholds of 25% sequence identity and 40% alignment coverage for both query and target. Using the alignment results, we created an undirected graph in NetworkX^82^ with each protein as a node, adding edges based on sequence similarity and clustering by connected components, resulting in 4,202 non-overlapping clusters. The NetworkX connected components method ensured no sequence similarity between proteins from different clusters.

We randomly partitioned the 4,202 clusters into training and validation sets in a 4:1 ratio, repeating this 10,000 times. To account for the uneven distribution of mutations, we ranked partitions by how closely their mutation counts matched the ideal 4:1 ratio, selecting the top five partitions. We verified that the pathogenic/benign ratio in these partitions matched the original dataset (see Table S4a).

To further evaluate the independence and diversity of partitions, we computed pairwise Jaccard similarity coefficients (Tables S4b, S4c). The similarity between training sets ranged from 0.63 to 0.70, indicating moderate overlap, while similarity between validation sets ranged from 0.05 to 0.17, confirming high distinctness and good complementarity.

We performed a grid search across these five partitions to identify the optimal hyperparameters, using the average area under the receiver operating characteristic curve (AU-ROC) as the performance metric. The final MutaPheno model was trained on the full training set using the selected parameters.

#### Statistical metrics and evaluation criteria

We evaluated model performance using a comprehensive set of metrics to capture different aspects of classification quality. These included: the area under the receiver operating characteristic curve (AUC-ROC), which assesses overall discriminative ability across thresholds; the area under the precision-recall curve (AUC-PR), which is particularly informative for imbalanced datasets; the maximum Matthews correlation coefficient (maximum MCC), representing the highest MCC achieved across thresholds and reflecting a balanced measure that incorporates all confusion matrix components; MCC computed using the classification labels generated by each method at its recommended threshold or default setting; F1-score, the harmonic mean of precision and recall; and precision and recall individually, where precision is the proportion of positive predictions that are correct, and recall is the proportion of actual positives correctly identified.

## Results and Discussion

To systematically investigate molecular-level characteristics of functional missense variants, we curated a comprehensive dataset spanning six major variant classes: pathogenic, benign, driver, passenger, recurrent somatic, and population variants. The dataset encompassed over 127,000 missense mutations affecting more than 15,000 proteins, each annotated with 34 molecular features describing structural and functional context, physicochemical changes, and broader protein-level properties. An overview of the study design and data composition is shown in Figure 1 and summarized in Table 1 and Supplementary Table S3. Mutation distributions across features are detailed in Figure 2 and Figure S1. As expected, features such as Core and ΔΔ*G*_*fold*_ covered a large number of mutations across variant classes, reflecting their extensive representation in protein structures. In contrast, features like SLiM, PNI-BS, and ΔΔ*G*_*bind*_ mapped to far fewer mutations, likely due to their localized nature and more limited structural coverage. These differences highlight variability in statistical power, which was accounted for in subsequent enrichment analyses.

**Figure 2.**
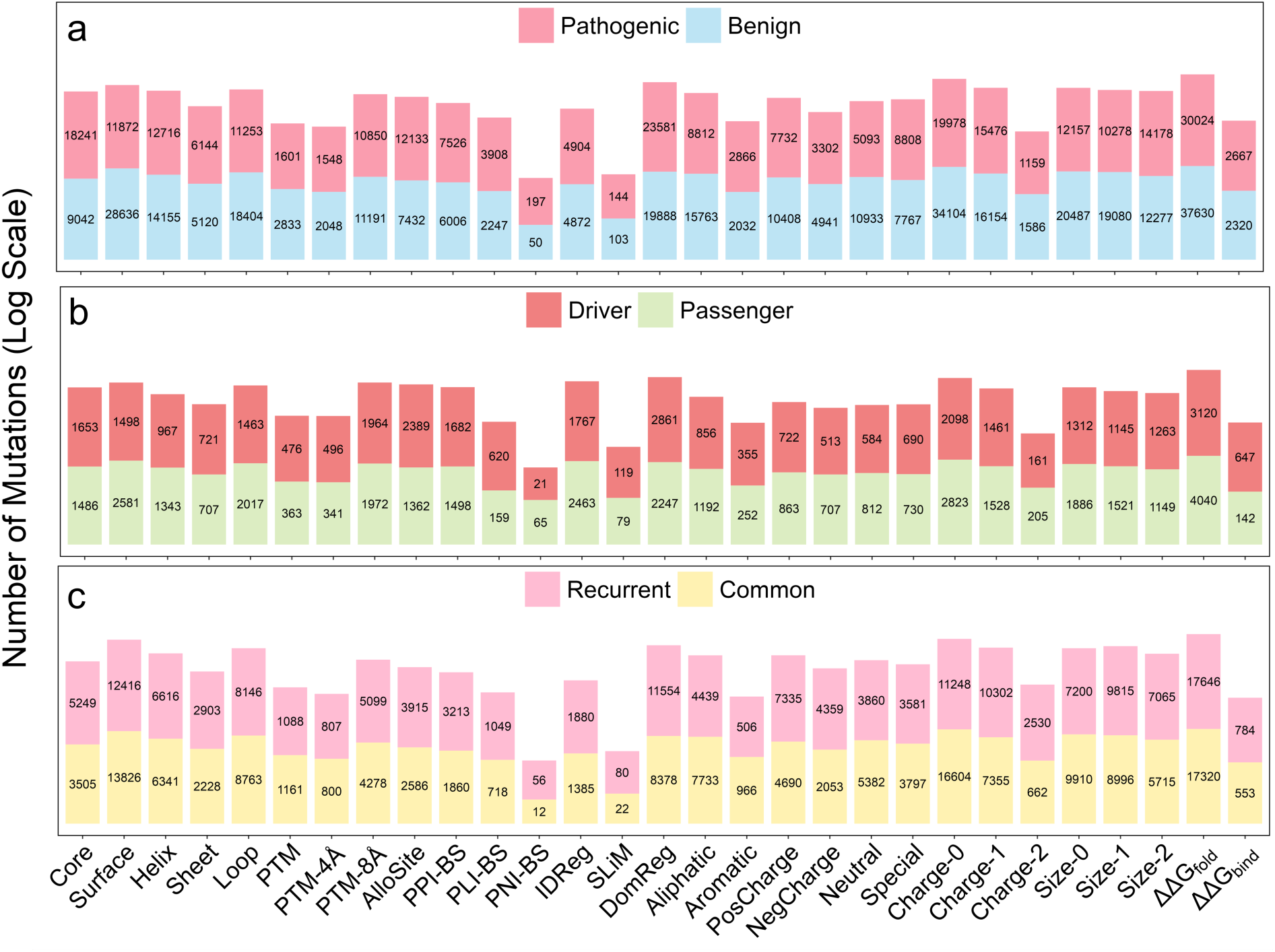
Mutation counts mapped to each of the 21 residue-level and 8 mutation-level features. (a) Pathogenic vs. benign mutations; (b) driver vs. passenger mutations; (c) recurrent somatic vs. common population mutations. Bars represent the total number of mutations falling within each annotated feature, with counts of each mutation class labeled directly on the bars. Mutation counts are plotted on a log₁₀ scale. Detailed statistics, including the number of mutations, unique mutated sites, and corresponding proteins, are provided in Data S2.

### Feature enrichment and functional association analysis

We first assessed 21 residue-level and six mutation-level features using odds ratio (OR) analysis across three binary classification tasks: Pathogenic vs. Benign, Driver vs. Passenger, and Recurrent vs. Common. The two remaining mutation-level features—folding stability and binding affinity changes—were evaluated based on continuous predicted ΔΔG values, as described in Section 4. For allosteric and binding sites, both experimental and computational annotations were available. We compared mutation enrichment (log₁₀OR) across tasks to assess consistency. Allosteric sites (Fig. S2a) and protein–ligand binding sites (Fig. S2c) showed consistent, significant enrichment, supporting their integration. Protein–protein binding sites (Fig. S2b) showed broadly consistent patterns except for divergence in Driver vs. Passenger comparisons; integration was retained to improve coverage. In contrast, protein–nucleic acid binding sites (Fig. S2d) showed inconsistent and often contradictory results, so only experimentally validated sites were used in downstream analyses. OR-based analyses across tasks (Figure 3) revealed distinct feature association patterns, allowing features to be grouped into four categories reflecting consistent or context-dependent associations with functional mutations.

**Figure 3.**
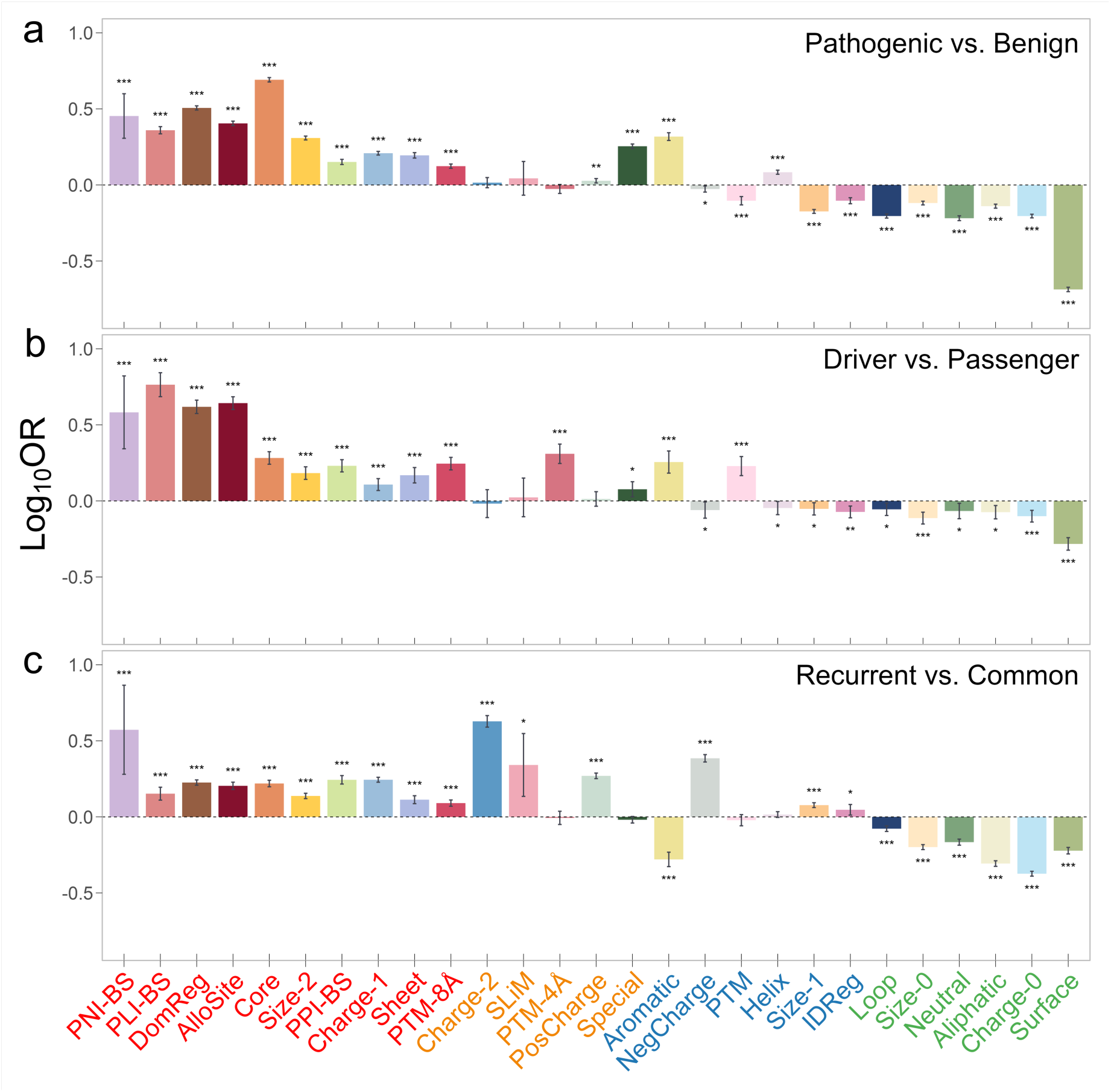
OR-based relative enrichment of residue- and mutation-level features across three mutation classification tasks. Panels show log₁₀ odds ratios (OR) comparing (a) pathogenic vs. benign, (b) driver vs. passenger, and (c) recurrent somatic vs. common population mutations for 21 residue-level and six mutation-level features. Error bars represent 95% confidence intervals. Significance was determined by Fisher’s exact test with Benjamini–Hochberg correction (*q < 0.05; **q < 0.001; ***q < 0.0001). Features were assigned to four color-coded categories—red, orange, blue, and green—based on the direction and significance of their ORs across all three comparisons. Within each category, features are ordered left to right by their mean log₁₀OR across the tasks. Underlying data for this figure are provided in Data S3.

#### Features with consistent associations with positive mutations

Ten features showed significant enrichment in positive mutations across all tasks (log₁₀OR > 0, q < 0.05), suggesting shared molecular signatures in both germline and somatic disease variants (red-labeled in Figure 3). Enriched features included functional sites such as PNI-BS, PLI-BS, DomReg, AlloSite, PPI-BS, and PTM-8Å, which are involved in transcription, catalysis, signaling, and post-translational regulation. Notably, PNI-BS showed consistent enrichment despite covering relatively few mutations, underscoring its functional relevance. Core and Sheet regions were also enriched, indicating that disruption of structural stability contributes to pathogenicity. Two mutation-level features, Size-2 and Charge-1, were consistently enriched, reflecting the impact of size and charge changes. These results highlight functional site disruption and structural destabilization as common mechanisms across disease-related mutations.

#### Features with context-dependent associations with positive mutations

Five features showed significant enrichment (log₁₀OR > 0, q < 0.05) in only a subset of classification tasks (orange-labeled in Figure 3). Among these, PTM-4Å exhibited the most pronounced differences between the Pathogenic vs. Benign and Driver vs. Passenger comparisons. PTM-4Å was enriched in somatic drivers, suggesting that mutations near post-translational modification sites may promote tumorigenesis by interfering with regulatory mechanisms.

#### Features with mixed-direction associations across mutation types

Six features showed significant but opposite enrichment patterns (q < 0.05) across different classification tasks (blue-labeled in Figure 3). Among them, PTM sites and Helix displayed opposing associations between germline and somatic contexts. PTM sites were enriched in driver mutations but depleted in pathogenic ones, suggesting a cancer-specific regulatory role. Conversely, Helix was enriched exclusively in pathogenic mutations, underscoring the importance of structural context in germline pathogenicity. These opposing trends indicate that the functional impact of molecular features can vary by mutation type, reinforcing the value of integrating diverse feature patterns into predictive models of mutation effects.

Compared to the Pathogenic vs. Benign and Driver vs. Passenger tasks, the Recurrent vs. Common classification showed distinct feature patterns for both orange- and blue-labeled features (Figure 3). This suggests that the forces driving mutation recurrence are different from those influencing pathogenicity or driver status. These differences highlight the importance of treating recurrence as a separate biological process, not just an extension of pathogenicity or driver classification.

#### Features with consistent associations with negative mutations

Six features showed significant and consistent negative associations across all tasks (log₁₀OR < 0, q < 0.05; green-labeled in Figure 3). Surface residues showed the strongest depletion, supporting the notion that solvent-exposed sites often tolerate mutations without functional consequences. Similarly, mutations with no change in charge (Charge-0) or size (Size-0) were underrepresented, likely due to their limited structural or physicochemical impact. Loops, aliphatic, and neutral residues were also consistently depleted, suggesting that flexible or chemically inert regions are less functionally constrained. Collectively, these patterns indicate that mutations with these features are more likely to be tolerated and functionally neutral across contexts.

### Integrative analysis of statistical associations and mutation enrichment

To complement odds ratio (OR) analysis, we applied Density Ratio (DR) metrics to assess the absolute enrichment of mutations across residue-level features. DR quantifies mutation density within specific regions relative to the global background, helping identify areas of potential functional or structural relevance. By integrating OR and DR (Figure 4), we identified features that are not only statistically associated with pathogenicity but also show localized enrichment, suggesting sites under selective pressure or biological vulnerability. DR profiles for all 21 residue-level features across six mutation datasets are provided in Figure S3.

**Figure 4.**
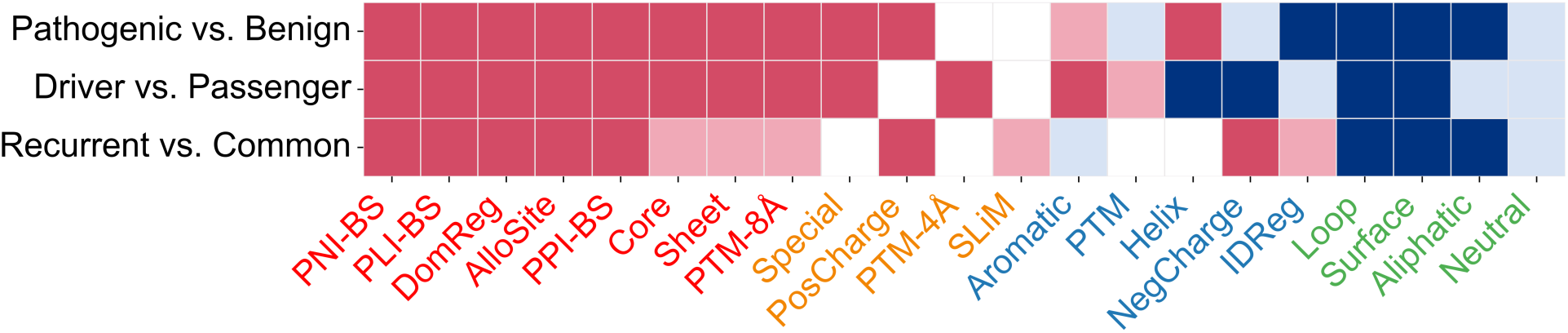
Integrated OR and DR enrichment for 21 residue-level features across mutation datasets. Each cell is color-coded by the feature’s OR and DR status: Dark red: log₁₀OR > 0 (q < 0.05) and log₁₀DR > 0 (q < 0.05) — significant association with positive mutations and actual enrichment. Pink: log₁₀OR > 0 (q < 0.05) but no DR enrichment (log₁₀DR ≤ 0 or q ≥ 0.05) — statistically linked to positive mutations without true enrichment. Dark blue: log₁₀OR < 0 (q < 0.05) and log₁₀DR > 0 (q < 0.05) — significant association with negative mutations and actual enrichment. Light blue: log₁₀OR < 0 (q < 0.05) but no DR enrichment (log₁₀DR ≤ 0 or q ≥ 0.05) — statistically linked to negative mutations without true enrichment. White: q ≥ 0.05 — no significant OR.

For example, nine features with consistent positive OR associations also showed significant DR enrichment in pathogenic and driver datasets (dark red in Figure 4; Fig. S3), indicating that these regions are both functionally critical and intrinsic mutational hotspots. Although Core, Sheet, and PTM-8Å significantly distinguished Recurrent from Common mutations, DR analysis revealed that this distinction was mainly driven by depletion of Common mutations rather than genuine enrichment of Recurrent ones (pink in Fig. 4; Fig. S3c). This supports the view that recurrent mutations are not necessarily functional drivers and that biologically important features may not always show enrichment for such mutations. For features with mixed-direction associations, DR analysis helped clarify apparent inconsistencies (blue-labeled in Fig. 4). For example, PTM sites showed significant OR associations but no significant DR enrichment in pathogenic and driver mutation categories (light blue and pink in Fig. 4), contrasting with prior reports of PTM enrichment in disease^83^. This underscores the added value of spatially extended features such as PTM-4Å and PTM-8Å, which may better capture the local structural and functional impact of PTMs.

### Stratified enrichment patterns of recurrent mutations improve driver prioritization

To further evaluate whether enrichment patterns of carefully selected molecular features can improve driver prioritization and help identify potential driver mutations, we systematically stratified recurrent somatic mutations by tumor-type recurrence patterns and gene-level categories and performed enrichment analyses. This stratification builds on previous studies proposing that mutations recurrent across multiple tumor types are more likely pan-cancer drivers under shared selective pressures, whereas single-tumor recurrent mutations may reflect cancer-type-specific drivers or background mutational processes^39, 84, 85^. Specifically, we distinguished single-tumor from multi-tumor recurrent mutations and further subdivided each group into four tiers by recurrence frequency to enable controlled comparisons and reduce confounding by recurrence level (Fig. 1a). Given the pivotal role of driver genes involved in key cancer pathways for tumorigenesis^5, 86^, we also classified genes into three cancer-associated subtypes, including a newly defined category of dual-function genes (CDG∩CPG), and a non-cancer pathogenic group. Mutation counts are summarized in Table 1, with feature distributions shown in Figure S1.

The log₁₀DR enrichment analysis of 21 residue-level features across 13 positive mutation categories (Fig. 5a) revealed that multi-tumor mutations, particularly those with higher recurrence (n > 2), exhibited DR patterns resembling known drivers, especially in functionally constrained features such as PLI-BS, DomReg, AlloSite, and SLiM. In contrast, mutations specific to a single tumor showed weak or no significant enrichment in the features commonly associated with drivers. At the gene level, mutations in dual-function genes (CDG∩CPG) showed the strongest and most consistent enrichment in functional features characteristic of drivers, underscoring their central role in oncogenesis. Mutations in CDG-only genes also showed notable but more moderate enrichment, whereas mutations in CPG and PG genes exhibited minimal or no enrichment, suggesting a weaker connection to key oncogenic processes. Hierarchical clustering based on log₁₀DR and q-value profiles (Fig. 5b) further illustrated these relationships. Pathogenic, driver, CDG∩CPG, and high-recurrence multi-tumor mutations clustered together, reflecting shared enrichment in functionally constrained regions and likely functional relevance. These results highlight that stratified enrichment patterns derived from our molecular features provide a robust framework for prioritizing protentional functional driver mutations, improving resolution beyond recurrence frequency alone.

**Figure 5.**
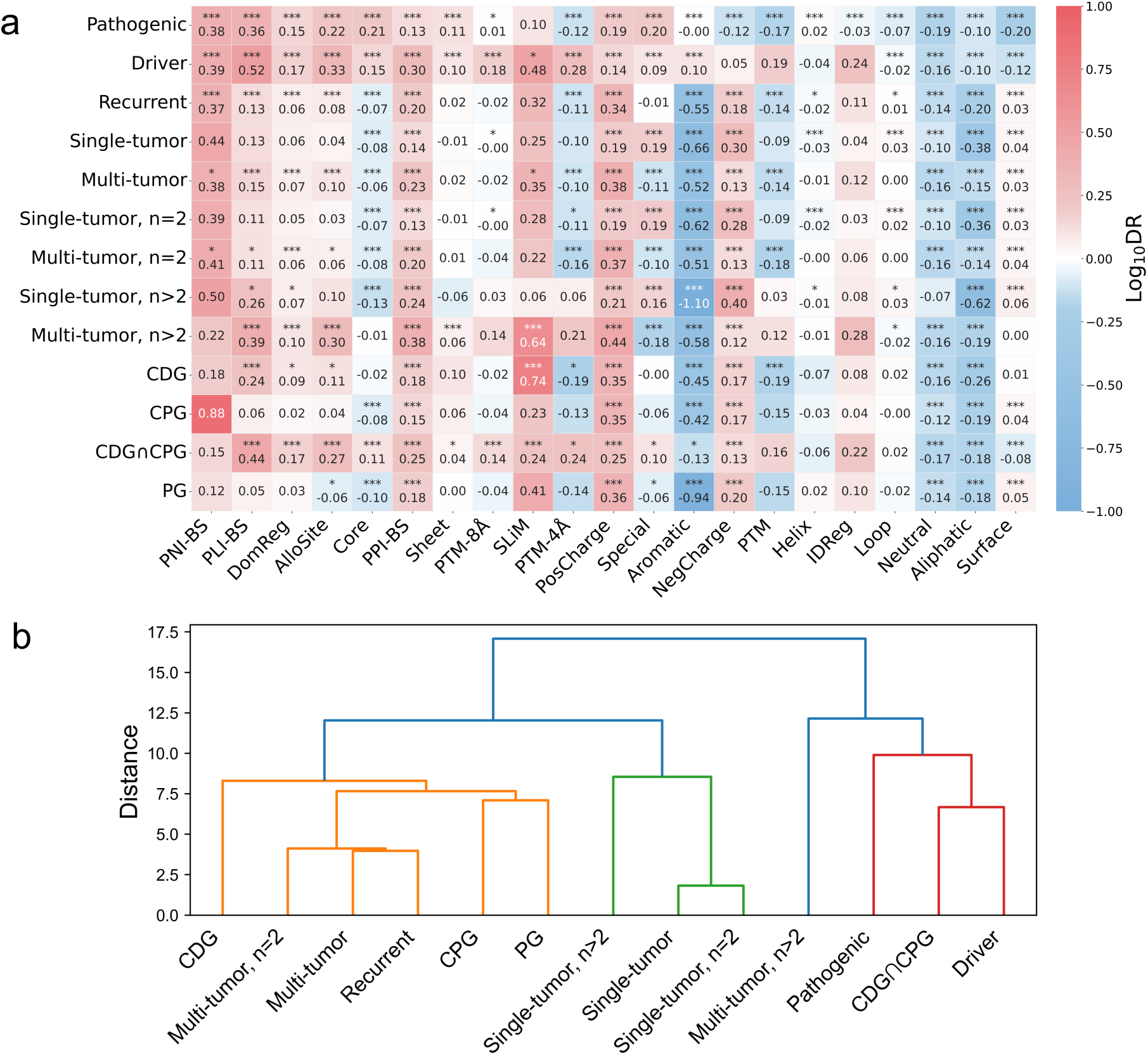
DR-based enrichment of 21 residue-level features across 13 mutation categories. (a) Heatmap of log₁₀DR values for each feature (columns) across mutation datasets (rows). Significance was assessed by comparing observed DR to a null distribution from 2,000 random simulations, with p-values adjusted via the BH method (*q < 0.05; **q < 0.001; ***q < 0.0001). (b) Hierarchical clustering of mutation categories based on their combined log₁₀DR and q-value profiles. The dendrogram groups categories by similarity in feature enrichment patterns. Underlying data are available in Data S4, and DR simulation results in Data S5.

### Biophysical disruption profiles distinguish driver-like mutations

The two biophysical mutation-level features—folding stability and binding affinity changes— were assessed using predicted continuous ΔΔG values. As shown in Fig. 6a, pathogenic mutations showed the strongest destabilization of folding (median ΔΔ*G*_*fold*_ significantly higher than benign variants), and drivers were similarly more destabilizing than passengers. Multi-tumor mutations with higher recurrence (n > 2) exhibited greater destabilization than Single-tumor events, consistent with stronger selection on pan-cancer drivers. At the gene level, CDG∩CPG mutations had the highest proportion of strongly destabilizing variants, with only this group’s mean ΔΔ*G*_*fold*_ significantly exceeding that of passengers.

**Figure 6.**
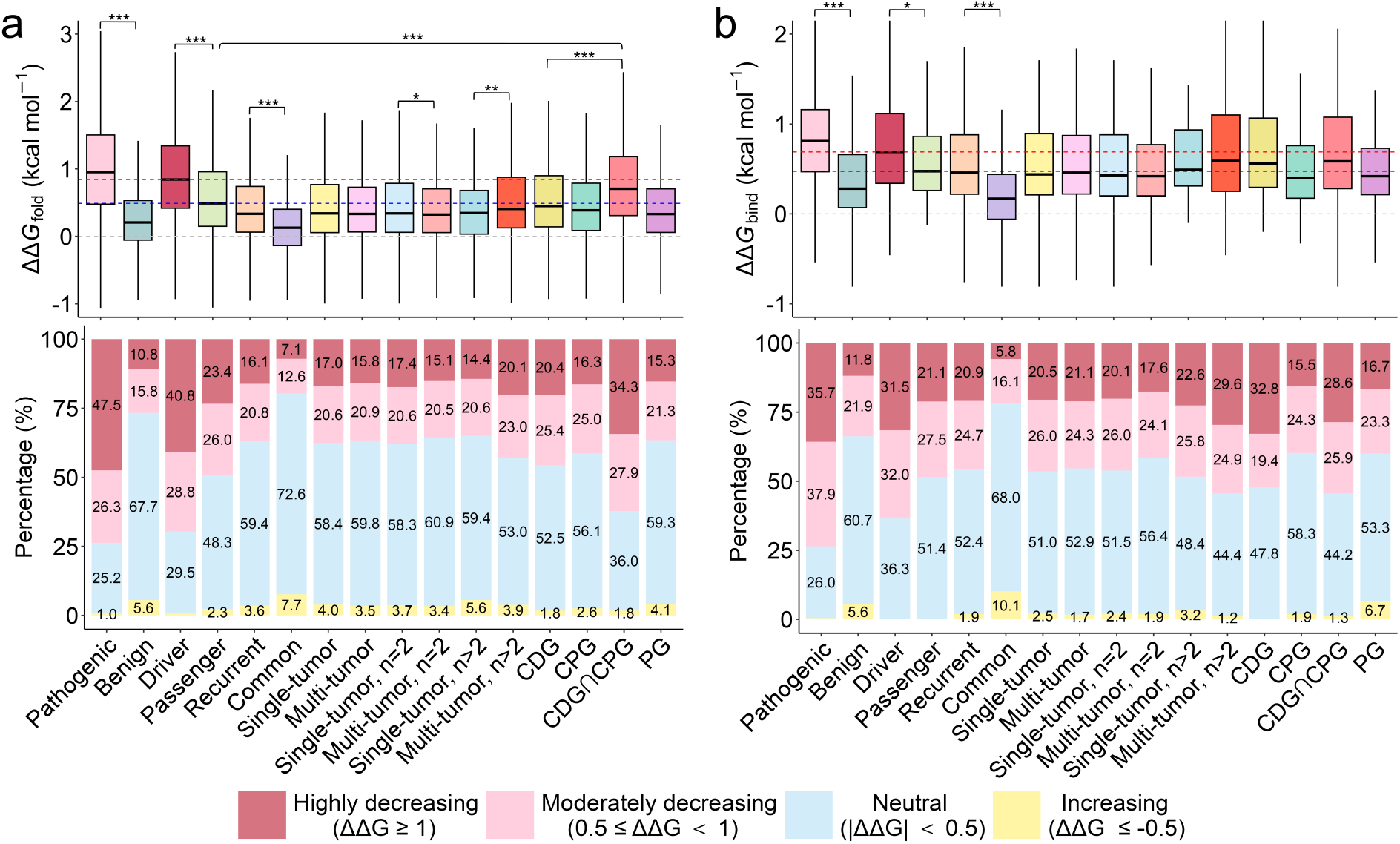
Predicted folding and binding perturbations across mutation categories. (a) Top: Box plots of predicted folding stability changes (ΔΔ*G*_*fold*_) for 16 mutation categories; bottom: stacked bars indicating the percentage of mutations in four ΔΔ*G*_*fold*_ categories: highly destabilizing, moderately destabilizing, neutral, and stabilizing. (b) Top: Box plots of predicted binding affinity changes (ΔΔ*G*_*bind*_) for the same categories; bottom: stacked bars of the four ΔΔ*G*_*bind*_ impact classes. Significance in the box plots was tested by Welch’s t-test with BH correction (*q < 0.05; **q < 0.001; ***q < 0.0001) across seven primary comparisons— pathogenic vs. benign; driver vs. passenger; recurrent vs. common; single- vs. multi-tumor (both n=2 and n>2 tiers); and CDG vs. CDG∩CPG—plus additional tests for groups exceeding the passenger median. The red, blue, and gray dashed lines on the box plots represent the median of driver mutations, the median of passenger mutations, and zero, respectively. Underlying data are in Data S6 (ΔΔ*G*_*fold*_) and Data S7 (ΔΔ*G*_*bind*_).

A parallel trend was observed for binding affinity: pathogenic and driver mutations caused greater disruption of protein–protein interactions than benign and passenger variants, with high-recurrence multi-tumor mutations again showing strong effects (Fig. 6b). CDG∩CPG and CDG groups were enriched for variants leading to substantial binding affinity loss, whereas CPG and PG groups showed minimal impacts. These findings demonstrate that folding and binding disruptions are robust indicators of mutational impact and highlight the value of ΔΔG-based features for prioritizing driver mutations in cancer^25^.

### MutaPheno and its feature-driven interpretability

MutaPheno is a random forest classifier trained on 67,791 high-confidence missense variants (30,113 pathogenic; 37,678 benign), all mapped to AlphaFold2-predicted structures. The model integrates 34 molecular features—21 residue-level, 8 mutation-level, and 5 aggregated protein-level annotations—each scaled by log₁₀OR enrichment or raw ΔΔG values. Hyperparameter tuning across five random training-validation partitions identified the optimal setting: 2300 trees with 8 features considered per split. Model performance was consistently strong (AUC-ROC 0.864 ± 0.009, MCC 0.580 ± 0.020; Table S5), demonstrating robustness across partitions. The final model was retrained on the full dataset using these parameters.

To elucidate the key determinants of MutaPheno’s predictions, we computed global SHAP (SHapley Additive exPlanations) values quantifying each feature’s average contribution (Fig. S4). Stability change emerged as the most influential factor (21.1% of total importance), followed by protein-level annotations such as KEGG pathway (13.6%), functional class (8.8%), and domain (8.6%). Among residue-level features, core/surface annotations (combined 14.1%) contributed most, while protein–protein binding sites (3.7%) and allosteric sites (3%) provided additional input. Sparsely represented features like SLiMs (0.04%) and PNI-BS (0.01%) contributed minimally— yet their nonzero importance underscores biological relevance, consistent with their enrichment patterns in Fig. 3. Among remaining mutation-level descriptors, large size changes (3.9%) and no charge change (3.1%) were most impactful. These results confirm that MutaPheno integrates structural and functional perturbations with multi-scale biological context, enabling robust and interpretable classification.

To further illustrate how MutaPheno generates interpretable predictions, we computed SHAP values for three well-characterized driver mutations (Fig. 7). For PIK3CA-E545K, top contributors included KEGG pathway membership, predicted disruption of p110α–p85α interactions (ΔΔ*G*_*bind*_ = 1.43 kcal/mol, based on PDB 4OVU), and allosteric site involvement (Fig. 7a). Consistent with prior studies reporting that E545K weakens this inhibitory interaction, mimicking activating phosphorylation, and driving constitutive PI3Kα activation^87^. KEGG pathways (e.g., ERBB, phosphatidylinositol, mTOR) are all tightly linked to PI3K/AKT signaling (Data S9). For TP53-R175H, the prediction was mainly driven by folding destabilization and location in the DNA-binding domain (Fig. 7b and Data S9), in line with experimental data showing that R175H disrupts the structural integrity of p53’s core domain, leading to loss of DNA binding^88^. For BRAF-V600E, SHAP drivers included strong ΔΔ*G*_*fold*_ and enrichment in MAPK pathway annotations (Fig. 7c and Data S9). This matches the known phosphomimetic role of V600E, which constitutively activates the MAPK/ERK signaling cascade^9, 89, 90^. These cases illustrate that MutaPheno not only delivers accurate predictions but also offers mechanistic insight, supporting its utility for hypothesis generation and experimental validation.

**Figure 7.**
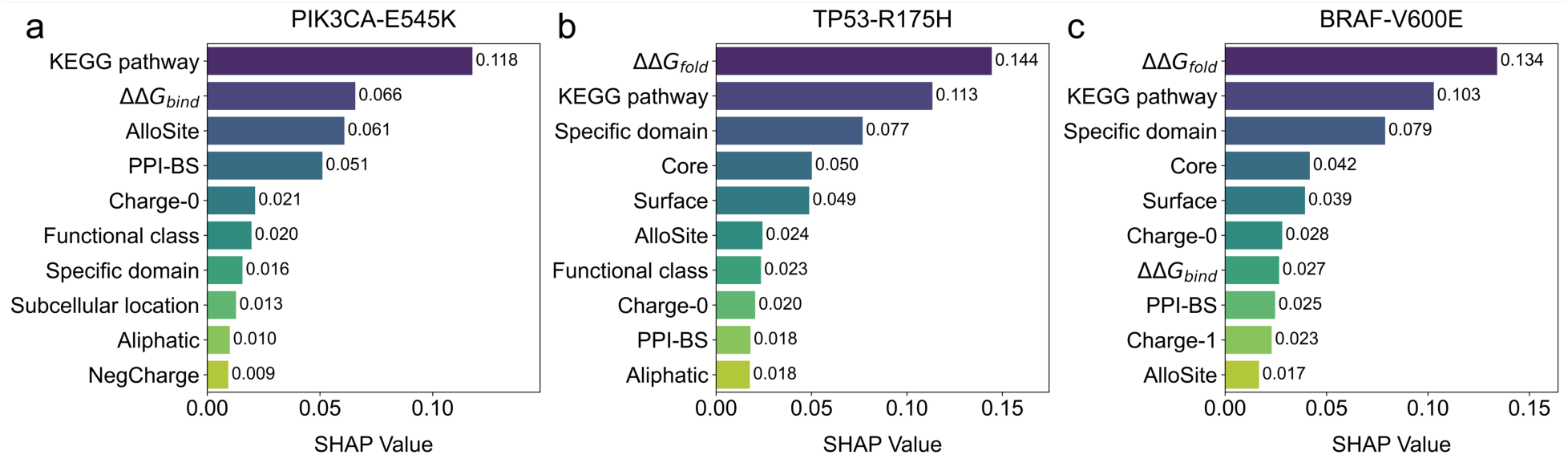
SHAP feature contributions for three representative driver mutations. Bars show the top 10 features ranked by absolute SHAP value for (a) PIK3CA E545K, (b) TP53 R175H, and (c) BRAF V600E. Higher SHAP values indicate greater influence on MutaPheno’s predicted pathogenic score for each specific mutation. Underlying data are available in Data S8.

### Benchmarking MutaPheno against existing pathogenicity and driver mutation prediction tools

We benchmarked MutaPheno on independent pathogenicity and driver mutation test sets to assess its predictive performance and generalizability. For pathogenicity, we evaluated 28,717 ClinVar-derived missense variants (12,539 pathogenic, 16,178 benign) and compared MutaPheno with 48 existing tools, including 47 models from dbNSFP4.9a^10, 11^ and CPT-1^77^. To ensure a fair comparison, we excluded 14 tools due to data leakage (e.g., MetaRNN, ClinPred, which were trained on ClinVar variants in the test set) or sparse coverage (<2% predictions returned, e.g., CADD). Among the remaining models, we focused on the top 10 performers: AlphaMissense, CPT-1, VEST4, DEOGEN2, MetaSVM, M-CAP, MVP, Eigen, MutFormer and our method MutaPheno. For direct comparison, we analyzed the common subset of 23,222 variants (10,485 pathogenic, 12,737 benign) with predictions from all 10 tools.

The performance of these 10 models using continuous probability values and discrete classification outputs is shown in Table 2 and Table S6, respectively. Based on probability scores, AlphaMissense achieved the highest predictive accuracy (AUC-ROC: 0.924; AUC-PR: 0.907), followed closely by CPT-1 and VEST4. MutaPheno ranked 7th in AUC-ROC (0.882) and achieving an AUC-PR of 0.867 (Fig. 8a). For models capable of providing classification outputs (Table S6), AlphaMissense again led the comparison (MCC: 0.700; F1-score: 0.830). MutaPheno ranked 5th in MCC (0.616) and achieved a strong F1-score (0.793), with balanced precision (0.775) and recall (0.812), confirming its solid classification performance relative to established tools. Notably, among these top-performing models, only MutaPheno provides interpretable predictions by linking molecular features to functional impact, offering added value for biological insight.

**Figure 8.**
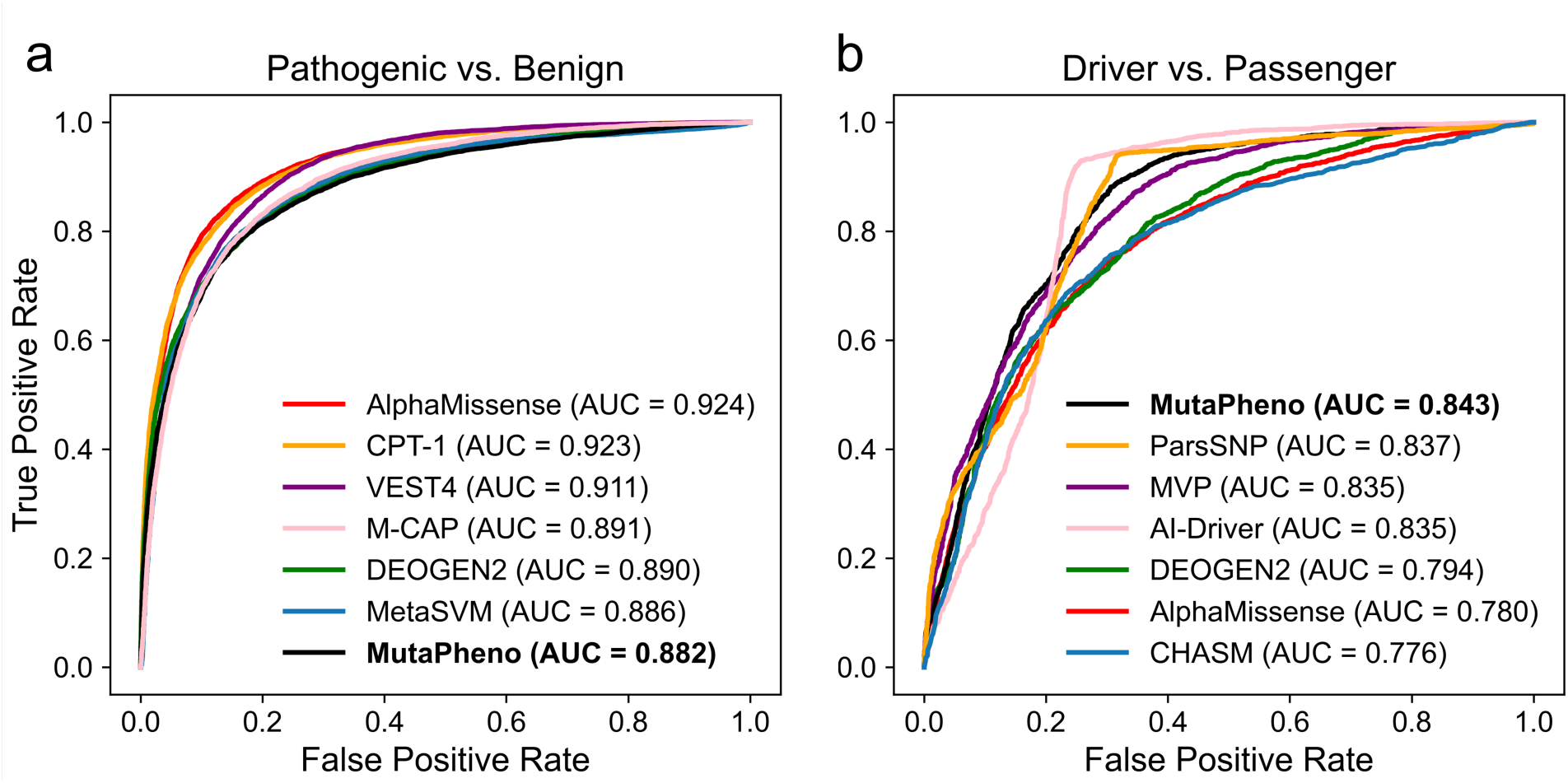
Comparative ROC analysis of predictive models across two mutation classification tasks. Receiver operating characteristic (ROC) curves for seven top-performing models on (a) pathogenic vs. benign and (b) driver vs. passenger variant classification. Area under the curve (AUC) values are shown in the legend for each model.

**Table 2.** Comparative performance of the top 10 pathogenicity prediction models on the intersected pathogenic mutation test set. Models are ranked in descending order of AUC-ROC. Variant-level prediction scores are provided in Data S9.

| **Model** | **AUC-ROC** | **AUC-PR** | **Maximum**<br>**MCC** | **Sensitivity**<br>**at 10% FPR** |
| --- | --- | --- | --- | --- |
| AlphaMissense | 0.924 | 0.907 | 0.703 | 0.792 |
| CPT-1 | 0.923 | 0.908 | 0.691 | 0.774 |
| VEST4 | 0.911 | 0.883 | 0.663 | 0.718 |
| M-CAP | 0.891 | 0.857 | 0.631 | 0.693 |
| DEOGEN2 | 0.890 | 0.879 | 0.623 | 0.700 |
| MetaSVM | 0.886 | 0.854 | 0.634 | 0.695 |
| **MutaPheno** | **0.882** | **0.867** | **0.622** | **0.683** |
| MVP | 0.882 | 0.856 | 0.612 | 0.670 |
| Eigen | 0.877 | 0.841 | 0.590 | 0.614 |
| MutFormer | 0.873 | 0.844 | 0.593 | 0.609 |

For driver prediction, we evaluated 6,322 somatic missense mutations (2,317 drivers and 4,005 passengers), comparing MutaPheno with both general pathogenicity predictors and well-established cancer-specific tools (CHASM, CHASMplus, ParsSNP, AI-Driver). MutaPheno achieved the best overall performance across the full dataset, outperforming all other models. For a controlled comparison, we analyzed 5,770 variants (2,018 drivers and 3,752 passengers) for which predictions were available from 12 representative models: the six top performers from the pathogenicity test set, MVP (the best general tool for driver prediction), the four cancer-specific tools, and MutaPheno. On this subset, MutaPheno achieved the highest AUC-ROC (0.843; Table 3, Fig. 8b), with ParsSNP and MVP also showing strong performance. Although AI-Driver achieved the highest maximum MCC, it had notably lower sensitivity at 10% FPR. Importantly, models that outperformed MutaPheno on pathogenicity showed substantially reduced accuracy for driver prediction (AUC-ROC < 0.800). In classification analyses (Table S7), MutaPheno achieved the second-highest MCC (0.530) and F1-score (0.707). As ParsSNP lacks direct classification outputs, applying its recommended cutoff (0.1) resulted in lower performance (MCC: 0.369).

**Table 3.**
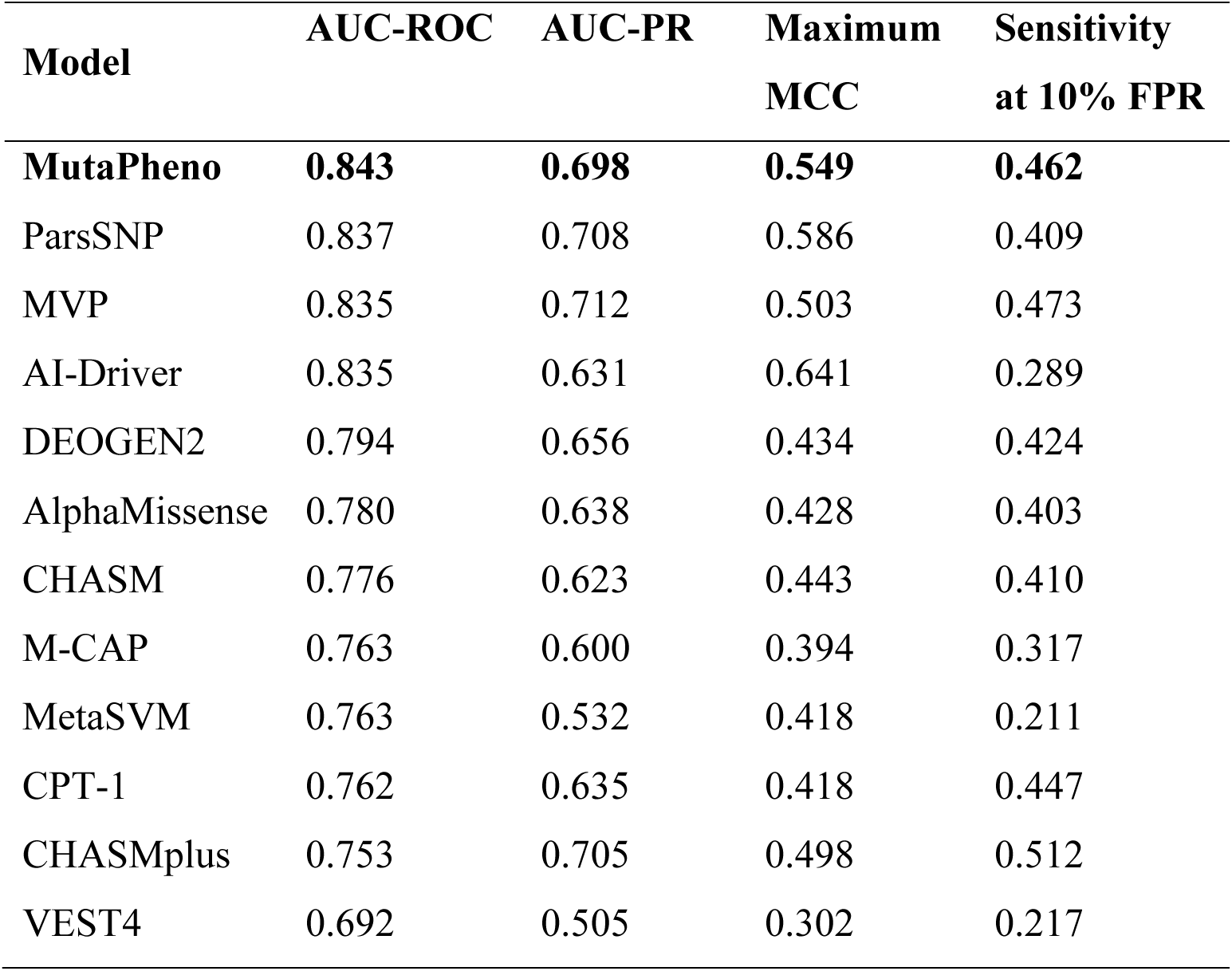
Comparative performance of the top 12 pathogenicity and cancer-specific prediction models on the intersected driver mutation test set. Models are ranked in descending order of AUC-ROC. Variant-level prediction scores are provided in Data S9.

| **Model** | **AUC-ROC** | **AUC-PR** | **Maximum**<br>**MCC** | **Sensitivity**<br>**at 10% FPR** |
| --- | --- | --- | --- | --- |
| **MutaPheno** | **0.843** | **0.698** | **0.549** | **0.462** |
| ParsSNP | 0.837 | 0.708 | 0.586 | 0.409 |
| MVP | 0.835 | 0.712 | 0.503 | 0.473 |
| AI-Driver | 0.835 | 0.631 | 0.641 | 0.289 |
| DEOGEN2 | 0.794 | 0.656 | 0.434 | 0.424 |
| AlphaMissense | 0.780 | 0.638 | 0.428 | 0.403 |
| CHASM | 0.776 | 0.623 | 0.443 | 0.410 |
| M-CAP | 0.763 | 0.600 | 0.394 | 0.317 |
| MetaSVM | 0.763 | 0.532 | 0.418 | 0.211 |
| CPT-1 | 0.762 | 0.635 | 0.418 | 0.447 |
| CHASMplus | 0.753 | 0.705 | 0.498 | 0.512 |
| VEST4 | 0.692 | 0.505 | 0.302 | 0.217 |

Notably, we excluded mutations overlapping with our training set to ensure an independent evaluation. However, ParsSNP, MVP, and AI-Driver were trained on data that partially overlapped with our test set, potentially inflating their reported performance. For example, AI-Driver’s training data included 1,665 driver and 532 passenger mutations present in our test set. ParsSNP employs a weakly supervised learning framework: during its unsupervised EM phase, it assigned driver probabilities to somatic mutations from TCGA, ICGC, and COSMIC; for example, 647 driver and 228 passenger mutations in our test set were present in the TCGA data used for label learning, and these probabilities were subsequently used as training labels.

To further evaluate our model’s predictive ability on unseen proteins, we performed leave-protein-out and leave-similar-protein-out testing. For pathogenicity, after removing variants from proteins in the training data, 338 variants remained (130 pathogenic, 208 benign). On this set, MutaPheno achieved an AUC-ROC of 0.841 and AUC-PR of 0.754 (Table S8a), demonstrating good discrimination on unseen proteins. For driver prediction, removing variants from proteins in the training set left fewer than 50 variants, so we adopted an alternative strategy: we retrained two models, MutaPheno* and MutaPheno^, with the same parameters. MutaPheno* (trained on proteins not overlapping with the test set, 63,839 mutations) achieved an AUC-ROC of 0.844, AUC-PR of 0.710, and MCC of 0.478 (Table S8b). MutaPheno^ (trained on proteins with <25% sequence similarity to the test set, 60,334 mutations) achieved an AUC-ROC of 0.841, AUC-PR of 0.699, and MCC of 0.475. These results highlight the model’s strong generalizability and robustness for unseen or distantly related proteins.

In this study, we systematically characterized the 34 molecular-level features of over 120,000 missense mutations spanning pathogenic, benign, driver, passenger, recurrent somatic, and population variants. Through comprehensive enrichment analyses, we identified distinct and shared molecular signatures across these mutation classes. Features related to functional sites (e.g., protein–protein, protein–ligand, nucleic acid binding sites; allosteric and domain regions) and structural integrity (e.g., core and sheet regions) consistently distinguished pathogenic and driver mutations from benign and passenger variants. These findings reinforce that key molecular interfaces and structural cores are vulnerable hotspots for functionally impactful mutations^26-28^. Our analysis of biophysical disruption (ΔΔ*G*_*fold*_ and ΔΔ*G*_*bind*_) further highlighted structural destabilization and interface perturbation as common hallmarks of disease-causing mutations across both germline and somatic contexts^25^. Features exhibiting mixed-direction associations across mutation types were rare; for example, only post-translational modification (PTM) sites and helix regions showed opposite enrichment trends between pathogenic and driver mutations, underscoring the overall consistency of molecular determinants underpinning these functional classes.

Our context-stratified analysis of recurrent somatic mutations provided additional insight into driver mutation prioritization. Multi-tumor recurrent mutations exhibited molecular profiles more similar to known driver mutations than single-tumor recurrent mutations, supporting the notion that pan-cancer recurrence reflects stronger selective pressure and functional relevance^39, 84, 85^. Mutations in dual-function genes (classified as both cancer drivers and pathway genes) showed the highest enrichment in functional features compared to other gene categories, highlighting their central role in oncogenesis, consistent with previous studies that emphasized the pivotal role of driver genes embedded in key cancer pathways in promoting tumorigenesis^5, 86^.

Building on these insights, we developed MutaPheno, an interpretable random forest classifier that predicts both pathogenic and cancer driver missense mutations based on 34 curated and computed molecular-level features spanning residue, mutation, and protein scales. By modeling molecular perturbations and biological context rather than directly mapping variants to labels, MutaPheno enhances both interpretability and generalizability across diverse contexts. A key finding is that despite being trained exclusively on pathogenic mutations, MutaPheno generalizes effectively to driver mutation prediction. This is attributable to the mechanistic convergence between pathogenic and driver mutations: although they arise in distinct clinical and evolutionary settings, both tend to disrupt similar molecular features. Our benchmarking confirmed this advantage: whereas many pathogenicity models (e.g., AlphaMissense, AUC-ROC decline from 0.924 to 0.780) exhibited marked performance drops on driver prediction tasks, MutaPheno’s AUC-ROC decreased only modestly (from 0.882 to 0.843). This robustness underscores the value of using biologically grounded, interpretable features and a random forest learning framework in achieving reliable transferability across variant types.

Despite these advances, certain limitations remain. The reliance on available structural and functional annotations constrains coverage for some proteins, particularly for protein–nucleic acid binding sites, short linear motifs, and protein–protein complex structures. Nonetheless, our work underscores the power of molecular-level features in mechanistically interpreting and predicting the impact of missense mutations. MutaPheno provides a robust and interpretable framework that lays the groundwork for future extensions. As functional datasets and structural resources continue to grow, integrating these additional data and expanding annotation coverage will further enhance the precision and utility of variant effect prediction in both precision medicine and cancer research.

## Data and Code Availability

The MutaPheno program is publicly available on GitHub at https://github.com/minghuilab/MutaPheno. The feature annotations for 16 mutation datasets can also be accessed on GitHub at the same link, with each dataset stored in a separate sheet. Each mutation is annotated with 34 molecular features. Additionally, the feature matrices and prediction results for the training set, as well as two independent test sets used in the development and evaluation of the MutaPheno model, can also be accessed on GitHub.

## Acknowledgments

This work was supported by the National Natural Science Foundation of China [32070665] and the Priority Academic Program Development of Jiangsu Higher Education Institutions. The funders had no role in study design, data collection and analysis, decision to publish, or preparation of the manuscript.

## Author Contributions

Conceptualization, M.L.; Methodology, Y.Y., W.S. and M.L.; Software, Y.Y., J.Z. and Y.L.; Validation, Y.Y. and M.L.; Formal Analysis, Y.Y. and M.L.; Investigation, Y.Y., W.S. and M.L.; Data Curation, Y.Y. and W.S.; Writing – Original Draft, Y.Y. and M.L.; Writing – Review & Editing, M.L.; Visualization, Y.Y. and M.L.; Supervision, M.L.; Project Administration, M.L.; Funding Acquisition, M.L.

## References

1. Zhao, F., Zheng, L., Goncearenco, A., Panchenko, A.R. & Li, M. Computational Approaches to Prioritize Cancer Driver Missense Mutations. Int. J. Mol. Sci. 19 (2018).

2. Stefl, S., Nishi, H., Petukh, M., Panchenko, A.R. & Alexov, E. Molecular mechanisms of disease-causing missense mutations. J. Mol. Biol. 425, 3919–3936 (2013).

3. Bunn, H.F. Pathogenesis and treatment of sickle cell disease. N. Engl. J. Med. 337, 762–769 (1997).

4. Hardy, J. & Selkoe, D.J. The amyloid hypothesis of Alzheimer’s disease: progress and problems on the road to therapeutics. Science 297, 353–356 (2002).

5. Vogelstein, B. et al. Cancer genome landscapes. Science 339, 1546–1558 (2013).

6. Stratton, M.R., Campbell, P.J. & Futreal, P.A. The cancer genome. Nature 458, 719–724 (2009).

7. Biankin, A.V. et al. Pancreatic cancer genomes reveal aberrations in axon guidance pathway genes. Nature 491, 399–405 (2012).

8. Chiang, Y.T. et al. The Function of the Mutant p53-R175H in Cancer. Cancers (Basel) 13 (2021).

9. Ascierto, P.A. et al. The role of BRAF V600 mutation in melanoma. J. Transl. Med. 10, 85 (2012).

10. Liu, X., Jian, X. & Boerwinkle, E. dbNSFP: a lightweight database of human nonsynonymous SNPs and their functional predictions. Hum. Mutat. 32, 894–899 (2011).

11. Liu, X., Li, C., Mou, C., Dong, Y. & Tu, Y. dbNSFP v4: a comprehensive database of transcript-specific functional predictions and annotations for human nonsynonymous and splice-site SNVs. Genome Med. 12, 103 (2020).

12. Ng, P.C. & Henikoff, S. SIFT: Predicting amino acid changes that affect protein function. Nucleic Acids Res. 31, 3812–3814 (2003).

13. Adzhubei, I.A. et al. A method and server for predicting damaging missense mutations. Nat. Methods 7, 248–249 (2010).

14. Ioannidis, N.M. et al. REVEL: An Ensemble Method for Predicting the Pathogenicity of Rare Missense Variants. Am. J. Hum. Genet. 99, 877–885 (2016).

15. Carter, H., Douville, C., Stenson, P.D., Cooper, D.N. & Karchin, R. Identifying Mendelian disease genes with the variant effect scoring tool. BMC Genomics 14 Suppl 3, S3 (2013).

16. Cheng, J. et al. Accurate proteome-wide missense variant effect prediction with AlphaMissense. Science 381, eadg7492 (2023).

17. Carter, H. et al. Cancer-specific high-throughput annotation of somatic mutations: computational prediction of driver missense mutations. Cancer Res. 69, 6660–6667 (2009).

18. Kumar, R.D., Swamidass, S.J. & Bose, R. Unsupervised detection of cancer driver mutations with parsimony-guided learning. Nat. Genet. 48, 1288–1294 (2016).

19. Tokheim, C. & Karchin, R. CHASMplus Reveals the Scope of Somatic Missense Mutations Driving Human Cancers. Cell Syst 9, 9–23.e28 (2019).

20. Wang, H., et al. AI-Driver: an ensemble method for identifying driver mutations in personal cancer genomes. NAR Genom Bioinform 2, lqaa084 (2020).

21. Ostroverkhova, D., Sheng, Y. & Panchenko, A. Are Next-Generation Pathogenicity Predictors Applicable to Cancer? J. Mol. Biol. 436, 168644 (2024).

22. Ostroverkhova, D., Przytycka, T.M. & Panchenko, A.R. Cancer driver mutations: predictions and reality. Trends Mol. Med. 29, 554–566 (2023).

23. Chen, H. et al. Comprehensive assessment of computational algorithms in predicting cancer driver mutations. Genome Biol. 21, 43 (2020).

24. Wierbowski, S.D., Fragoza, R., Liang, S. & Yu, H. Extracting Complementary Insights from Molecular Phenotypes for Prioritization of Disease-Associated Mutations. Curr Opin Syst Biol 11, 107–116 (2018).

25. Li, M. et al. Balancing Protein Stability and Activity in Cancer: A New Approach for Identifying Driver Mutations Affecting CBL Ubiquitin Ligase Activation. Cancer Res. 76, 561–571 (2016).

26. Iqbal, S. et al. Comprehensive characterization of amino acid positions in protein structures reveals molecular effect of missense variants. Proc. Natl. Acad. Sci. U. S. A. 117, 28201–28211 (2020).

27. Cheng, F. et al. Comprehensive characterization of protein-protein interactions perturbed by disease mutations. Nat. Genet. 53, 342–353 (2021).

28. Laddach, A., Ng, J.C.F. & Fraternali, F. Pathogenic missense protein variants affect different functional pathways and proteomic features than healthy population variants. PLoS Biol. 19, e3001207 (2021).

29. Gerasimavicius, L., Livesey, B.J. & Marsh, J.A. Loss-of-function, gain-of-function and dominant-negative mutations have profoundly different effects on protein structure. Nat Commun 13, 3895 (2022).

30. Pires, D.E., Chen, J., Blundell, T.L. & Ascher, D.B. In silico functional dissection of saturation mutagenesis: Interpreting the relationship between phenotypes and changes in protein stability, interactions and activity. Sci. Rep. 6, 19848 (2016).

31. Landrum, M.J. et al. ClinVar: public archive of relationships among sequence variation and human phenotype. Nucleic Acids Res. 42, D980–985 (2014).

32. UniProt: the Universal Protein Knowledgebase in 2023. Nucleic Acids Res. 51, D523-d531 (2023).

33. Sevim Bayrak, C., et al. Identification of discriminative gene-level and protein-level features associated with pathogenic gain-of-function and loss-of-function variants. Am. J. Hum. Genet. 108, 2301–2318 (2021).

34. Stenson, P.D. et al. The Human Gene Mutation Database (HGMD(®)): optimizing its use in a clinical diagnostic or research setting. Hum. Genet. 139, 1197–1207 (2020).

35. Chakravarty, D. et al. OncoKB: A Precision Oncology Knowledge Base. JCO Precis Oncol 2017 (2017).

36. Tamborero, D. et al. Cancer Genome Interpreter annotates the biological and clinical relevance of tumor alterations. Genome Med. 10, 25 (2018).

37. Brown, A.L., Li, M., Goncearenco, A. & Panchenko, A.R. Finding driver mutations in cancer: Elucidating the role of background mutational processes. PLoS Comput. Biol. 15, e1006981 (2019).

38. Ellrott, K. et al. Scalable Open Science Approach for Mutation Calling of Tumor Exomes Using Multiple Genomic Pipelines. Cell Syst 6, 271–281.e277 (2018).

39. Bailey, M.H. et al. Comprehensive Characterization of Cancer Driver Genes and Mutations. Cell 173, 371–385.e318 (2018).

40. Lek, M. et al. Analysis of protein-coding genetic variation in 60,706 humans. Nature 536, 285–291 (2016).

41. Chang, M.T. et al. Identifying recurrent mutations in cancer reveals widespread lineage diversity and mutational specificity. Nat. Biotechnol. 34, 155–163 (2016).

42. Nightingale, A. et al. The Proteins API: accessing key integrated protein and genome information. Nucleic Acids Res. 45, W539–w544 (2017).

43. Zaru, R. & Orchard, S. UniProt Tools: BLAST, Align, Peptide Search, and ID Mapping. Curr Protoc 3, e697 (2023).

44. Sievers, F. et al. Fast, scalable generation of high-quality protein multiple sequence alignments using Clustal Omega. Mol. Syst. Biol. 7, 539 (2011).

45. Martínez-Jiménez, F. et al. A compendium of mutational cancer driver genes. Nat. Rev. Cancer 20, 555–572 (2020).

46. Wang, T. et al. OncoVar: an integrated database and analysis platform for oncogenic driver variants in cancers. Nucleic Acids Res. 49, D1289–d1301 (2021).

47. Repana, D. et al. The Network of Cancer Genes (NCG): a comprehensive catalogue of known and candidate cancer genes from cancer sequencing screens. Genome Biol. 20, 1 (2019).

48. Sondka, Z. et al. The COSMIC Cancer Gene Census: describing genetic dysfunction across all human cancers. Nat. Rev. Cancer 18, 696–705 (2018).

49. Rehm, H.L. et al. ClinGen--the Clinical Genome Resource. N. Engl. J. Med. 372, 2235–2242 (2015).

50. Subramanian, A. et al. Gene set enrichment analysis: a knowledge-based approach for interpreting genome-wide expression profiles. Proc. Natl. Acad. Sci. U. S. A. 102, 15545–15550 (2005).

51. Raju, R. et al. NetSlim: high-confidence curated signaling maps. Database (Oxford) 2011, bar032 (2011).

52. Kabsch, W. & Sander, C. Dictionary of protein secondary structure: pattern recognition of hydrogen-bonded and geometrical features. Biopolymers 22, 2577–2637 (1983).

53. Liu, X. et al. Unraveling allosteric landscapes of allosterome with ASD. Nucleic Acids Res. 48, D394–d401 (2020).

54. Song, K. et al. Improved Method for the Identification and Validation of Allosteric Sites. J. Chem. Inf. Model. 57, 2358–2363 (2017).

55. Meyer, M.J. et al. Interactome INSIDER: a structural interactome browser for genomic studies. Nat. Methods 15, 107–114 (2018).

56. Krivák, R. & Hoksza, D. P2Rank: machine learning based tool for rapid and accurate prediction of ligand binding sites from protein structure. J. Cheminform. 10, 39 (2018).

57. Xia, Y., Xia, C.Q., Pan, X. & Shen, H.B. GraphBind: protein structural context embedded rules learned by hierarchical graph neural networks for recognizing nucleic-acid-binding residues. Nucleic Acids Res. 49, e51 (2021).

58. Tak Leung, R.W., Jiang, X., Chu, K.H. & Qin, J. ENPD - A Database of Eukaryotic Nucleic Acid Binding Proteins: Linking Gene Regulations to Proteins. Nucleic Acids Res. 47, D322–d329 (2019).

59. Liao, J.Y. et al. EuRBPDB: a comprehensive resource for annotation, functional and oncological investigation of eukaryotic RNA binding proteins (RBPs). Nucleic Acids Res. 48, D307–d313 (2020).

60. Hornbeck, P.V., et al. PhosphoSitePlus, 2014: mutations, PTMs and recalibrations. Nucleic Acids Res. 43, D512–520 (2015).

61. Chung, C.R., et al. dbPTM 2025 update: comprehensive integration of PTMs and proteomic data for advanced insights into cancer research. Nucleic Acids Res. 53, D377–d386 (2025).

62. Piovesan, D., et al. MOBIDB in 2025: integrating ensemble properties and function annotations for intrinsically disordered proteins. Nucleic Acids Res. 53, D495–d503 (2025).

63. Aspromonte, M.C., et al. DisProt in 2024: improving function annotation of intrinsically disordered proteins. Nucleic Acids Res. 52, D434–d441 (2024).

64. Kumar, M. et al. ELM-the Eukaryotic Linear Motif resource-2024 update. Nucleic Acids Res. 52, D442–d455 (2024).

65. Blum, M. et al. InterPro: the protein sequence classification resource in 2025. Nucleic Acids Res. 53, D444–d456 (2025).

66. Chen, Y. et al. PremPS: Predicting the impact of missense mutations on protein stability. PLoS Comput. Biol. 16, e1008543 (2020).

67. Zhang, N. et al. MutaBind2: Predicting the Impacts of Single and Multiple Mutations on Protein-Protein Interactions. iScience 23, 100939 (2020).

68. Berman, H.M. et al. The Protein Data Bank. Nucleic Acids Res. 28, 235–242 (2000).

69. Jumper, J. et al. Highly accurate protein structure prediction with AlphaFold. Nature 596, 583–589 (2021).

70. Varadi, M. et al. AlphaFold Protein Structure Database in 2024: providing structure coverage for over 214 million protein sequences. Nucleic Acids Res. 52, D368–d375 (2024).

71. Virtanen, P. et al. SciPy 1.0: fundamental algorithms for scientific computing in Python. Nat. Methods 17, 261–272 (2020).

72. Benjamini, Y. & Hochberg, Y. Controlling the False Discovery Rate: A Practical and Powerful Approach to Multiple Testing. Journal of the Royal Statistical Society: Series B (Methodological) 57, 289–300 (1995).

73. Altman, D.G. & Bland, J.M. Interaction revisited: the difference between two estimates. BMJ 326, 219 (2003).

74. Deák, G. & Cook, A.G. Missense Variants Reveal Functional Insights Into the Human ARID Family of Gene Regulators. J. Mol. Biol. 434, 167529 (2022).

75. Thomas, P.D. et al. PANTHER: Making genome-scale phylogenetics accessible to all. Protein Sci. 31, 8–22 (2022).

76. Thul, P.J. et al. A subcellular map of the human proteome. Science 356 (2017).

77. Jagota, M. et al. Cross-protein transfer learning substantially improves disease variant prediction. Genome Biol. 24, 182 (2023).

78. Li, M., Simonetti, F.L., Goncearenco, A. & Panchenko, A.R. MutaBind estimates and interprets the effects of sequence variants on protein-protein interactions. Nucleic Acids Res. 44, W494–501 (2016).

79. Sun, T., Chen, Y., Wen, Y., Zhu, Z. & Li, M. PremPLI: a machine learning model for predicting the effects of missense mutations on protein-ligand interactions. Commun Biol 4, 1311 (2021).

80. Zheng, F., Jiang, X., Wen, Y., Yang, Y. & Li, M. Systematic investigation of machine learning on limited data: A study on predicting protein-protein binding strength. Comput Struct Biotechnol J 23, 460–472 (2024).

81. Steinegger, M. & Söding, J. MMseqs2 enables sensitive protein sequence searching for the analysis of massive data sets. Nat. Biotechnol. 35, 1026–1028 (2017).

82. Hagberg, A.A., Schult, D.A., Swart, P. & Hagberg, J. Exploring Network Structure, Dynamics, and Function using NetworkX. Proceedings of the Python in Science Conference (2008).

83. Reimand, J., Wagih, O. & Bader, G.D. Evolutionary constraint and disease associations of post-translational modification sites in human genomes. PLoS Genet. 11, e1004919 (2015).

84. Priestley, P. et al. Pan-cancer whole-genome analyses of metastatic solid tumours. Nature 575, 210–216 (2019).

85. Hess, J.M. et al. Passenger Hotspot Mutations in Cancer. Cancer Cell 36, 288–301.e214 (2019).

86. The ICGC/TCGA Pan-Cancer Analysis of Whole Genomes Consortium. Pan-cancer analysis of whole genomes. Nature 578, 82-93 (2020).

87. Leontiadou, H., Galdadas, I., Athanasiou, C. & Cournia, Z. Insights into the mechanism of the PIK3CA E545K activating mutation using MD simulations. Sci. Rep. 8, 15544 (2018).

88. Chen, X. et al. Mutant p53 in cancer: from molecular mechanism to therapeutic modulation. Cell Death Dis. 13, 974 (2022).

89. Roviello, G. et al. Advances in anti-BRAF therapies for lung cancer. Invest. New Drugs 39, 879–890 (2021).

90. Kiel, C., Benisty, H., Lloréns-Rico, V. & Serrano, L. The yin-yang of kinase activation and unfolding explains the peculiarity of Val600 in the activation segment of BRAF. Elife 5, e12814 (2016).

